# A systems biology approach to evaluate potential probiotic candidates for women’s vaginal health

**DOI:** 10.1101/2025.06.16.659967

**Authors:** Emma M. Glass, Glynis L. Kolling, Jason A. Papin

## Abstract

Probiotic supplements are marketed for diverse health benefits, yet species inclusion often lacks functional rationale. We surveyed 352 U.S. probiotic products and found 36 unique microbial species, with most supplements containing only one species and no clear link between species and intended health benefit. To evaluate probiotic function, we developed CoPaPro, a collection of 1,012 genome-scale metabolic models spanning commensal, pathogenic, and probiotic bacteria. Flux balance analysis revealed that current probiotic species fail to capture the metabolic diversity of native commensals. Focusing on vaginal health, we identified commensals with metabolic profiles overlapping *Gardnerella vaginalis*, a key pathobiont. *In vitro* spent media assays using 11 vaginal isolates showed variable inhibition of *G. vaginalis*, primarily driven by D-lactic acid production rather than metabolic similarity. Several non-*Lactobacillus* species produced inhibitory levels of D-lactate. These findings highlight the need for function-based probiotic design and demonstrate a scalable framework integrating metabolic modeling with experimental validation.

## Introduction

Probiotics are defined as live microorganisms that when administered in adequate amounts confer a health benefit to the host (*1*). Many fermented foods containing probiotic lactic acid bacteria (*Lactobacilllus, Bacillus,* and *Bifidobacterium*) have been consumed for centuries, including kimchi, sauerkraut, and yogurt, among others (*2*). Around the year 2000 (S1) research on probiotics and probiotic rich foods began to increase exponentially resulting in a plethora of studies touting the extensive health benefits of probiotics including obesity prevention, improved gut-health, reduced risk of metabolic disorders, modulation of fecal microbiota, improvement of liver metabolism, and reducing cholesterol (*3–6*).

The commercial probiotics market was estimated at $70 billion in 2022 and is expected to hit $374 billion by 2034 (*7*). Despite the substantial probiotics market size and rapid growth, there have only been two probiotic products that have been categorized as therapeutic and approved by the FDA in the United States (biologics VOWST and REBYOTA for treating recurrent *Clostridioides difficile* infection) (*8*, *9*). The rest of the probiotic market is occupied by probiotic supplements that are marketed for specific uses such as improving gut health, vaginal health, and even brain health. However, these claims are often not backed by substantial *in vitro* scientific research, preclinical, or clinical studies, due to the low level of necessary FDA regulation of these products (*10*, *11*). This observation highlights the gap in understanding how species included in probiotic supplements can functionally and mechanistically support the targeted use case. It is necessary that we gain a deeper understanding of the range of metabolic functions across the species included in current probiotic supplements to develop more intentional and rationally designed probiotics for specific use cases.

One use case that is being increasingly targeted by probiotic companies is women’s health, specifically vaginal microbiome health (*12–15*). The vaginal microbiome is essential for maintaining vaginal health and protecting against dysbiosis and opportunistic pathogen proliferation (*16*). Generally, a vaginal microbiome that is considered a healthy state in reproductive age women is dominated by *Lactobacillus*, producing lactic acid to maintain an acidic vaginal pH (*16*). In a shift to a dysbiotic state, populations of non-*Lactobacillus* commensal vaginal bacteria take hold, resulting in an increase in species diversity in the microbiome and a higher pH (*17*). This shift leaves the vaginal microbiome susceptible to bacterial vaginosis (BV), a dysbiotic vaginal state often characterized by the presence of *Gardnerella vaginalis* (*18*, *19*). However, there are also populations of women with naturally more diverse vaginal microbiomes that do not present with BV-related symptoms. It has been shown that the functional niches of these diverse vaginal microbiomes overlap with both *G. vaginalis* and *Lactobacillus sp.*, resulting in prevention of both *G. vaginalis* proliferation and *Lactobacillus* dominance (*20*). BV is typically treated in the clinic with either oral antibiotics or vaginal antibiotic suppositories; however, there is a high rate of BV recurrence reported in vulnerable populations (*21–23*). Additionally, the misuse, overuse, and off-target effects of broad-spectrum antibiotics in clinical settings is rapidly leading to the rise of antimicrobial resistant bacteria, resulting in infections that are becoming increasingly difficult to treat (*24*). This heterogeneity in vaginal microbiome composition across women of varying demographics and high rates of BV recurrence after antibiotic treatments highlights the need for something other than a one-size-fits all approach to treat BV.

One promising avenue to explore is probiotic therapeutics and preventative supplements. Interestingly, there is research showing that oral or vaginal suppositories of *Lactobacillus* species can restore equilibrium in the vaginal microbiome (*25*, *26*). We performed cross-cohort meta-analysis of 32 independent research studies that treated recurrent BV with probiotics, antibiotics, or probiotic/antibiotic combination therapies (S2, S3). Across these 32 studies, we see no statistical difference in clinical outcomes of patients treated with oral antibiotics, intravaginal probiotics, antibiotics, or probiotic/antibiotic combination therapies in terms of BV recurrence. We also observed a large variation in effectiveness at preventing BV recurrence across groups. This underscores the fact that probiotics are a promising avenue for preventing BV recurrence, as they seem to be just as effective as antibiotic therapies. Additionally, because of the large variation in recurrence across treatment modalities, there is an opportunity to mechanistically and rationally design a novel and more effective probiotic supplement to support vaginal microbiome health by preventing BV and *Gardnerella vaginalis* colonization.

In this study, we perform an extensive survey of probiotic supplement products available at the top three pharmacies in the United States to gain a more comprehensive understanding of the probiotic market landscape. Additionally, we use a metabolic network modeling approach to better understand the mechanistic landscape of bacterial species used in probiotic supplements. Genome-scale metabolic network reconstructions (GENREs) are powerful modeling tools that can provide metabolic context to ‘omics data and simulate metabolic phenotypes of bacteria at the strain-level (*27*). GENREs enable us to explore the functional similarities and differences between probiotic, commensal, and pathogenic bacteria through species-specific network model generation and constraint-based analysis (COBRA) (*28*). Through network model simulation, we identify unique metabolic signatures of probiotic species and determine that there are functional gaps in the landscape of current probiotic species. Secondly, we use a combination of computational and *in vitro* approaches to identify existing commensal members of the vaginal microbiome that could be candidates for a novel rationally designed probiotic supplement for preventing vaginal dysbiosis. To accomplish this, we leverage GENREs to better understand the similarities and differences in metabolic function of commensal vaginal species. Secondly, we perform *in vitro* spent media experiments to determine which commensal vaginal bacteria can inhibit *G. vaginalis* growth.

## Results

### Survey of 352 over the counter probiotic supplements

To gain a deeper understanding of the landscape of over-the-counter probiotic supplements and the diversity of bacterial species they contain, we performed a survey of all over-the-counter probiotic supplements available at the top three pharmacies in the United States; CVS, Walgreens, and Walmart (9,600, 9,323, and 4,865 store locations respectively) (*29*). Across these pharmacies there were 352 unique probiotic supplement products available for purchase. Across these 352 supplements, there were 36 unique probiotic species used (35 bacterial species; 1 fungal species, *Saccharomyces boulardii*), and 70 distinct brand names (Figure 1 A).

**Figure 1.**
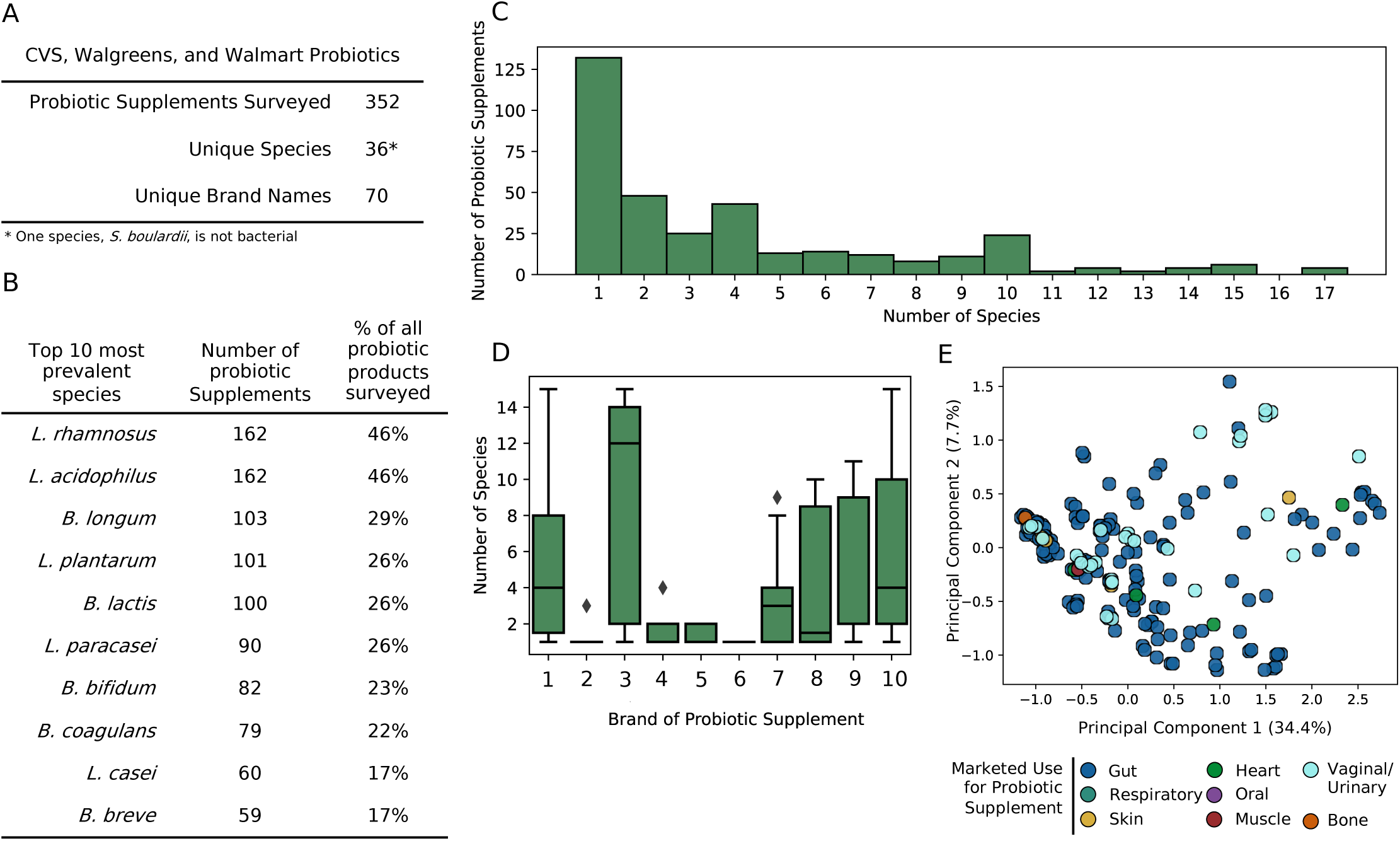
Survey of over the counter probiotic supplements. A) Description of products surveyed across CVS, Walgreens and Walmart. B) Top 10 most prevalent probiotic species across surveyed supplements. C) Histogram of the number of probiotic supplements containing a certain number of probiotic species. D) Box and whisker plot of the number of species contained in a probiotic supplement by brand-name. Only brand-names containing three or more unique products were displayed on this plot. E) PCA plot of the species profiles of each probiotic supplement. Colors of points represent the marketed use for each probiotic supplement.

The 10 most prevalent species of bacteria found across the 352 probiotic supplements are detailed in Figure 1 B. The top two species *L. rhamnosus* and *L. acidophilus* were present in 46% of all probiotic supplements. Both species are lactic acid-producing bacteria that are also found naturally in fermented dairy products like yogurt and cheese. Additionally, *B. coagulans* was present in 22% of probiotic supplements, most often appearing in probiotic gummies. *B. coagulans* spores are added to gummy-based probiotics due to their high survival rates during higher-temperature processing steps required for creating gummies (*30*). Other species like *Lactobacillus* and *Bifidobacteria* are typically lyophilized and encapsulated (*31*).

Additionally, we recorded the number of species included in each supplement (Figure 1 C) noting over half of the probiotic supplements contain one probiotic species. We observed over half of the probiotic supplements only contained one probiotic species. This result was surprising, because it suggests that the administration of one probiotic species can have a significant impact on health. Studies have shown that multi-strain probiotics are more effective at promoting biological activities and displacing pathogens due to the synergy amongst probiotic strains (*32*). This observation suggests there could be a benefit to taking a multi species probiotic, despite most over-the-counter probiotic supplements only containing one species. However, there are several supplements that contain more than five unique species and even several that contain 17 unique species.

Because of the variation in number of unique probiotic species included in a supplement, we explored if the number of unique species was linked with brand-name (Figure 1 D, Brand-names have been de-identified). We observed that name brand #3 probiotics have a median of 12 species in each product. On the other hand, name brand #6 probiotics have a median of 1 species in each product. Additionally, some brand names have a significant spread in the number of probiotic species included across products. For example, name brand #1, name brand #3, and name brand #10 probiotics all range from having one species to 15 species.

Because of the apparent inconsistency in the number and type of species used in probiotic supplements, we wanted to determine whether there is a link between the species included in a supplement, and the marketed use for the probiotic. We recorded the targeted use for supplement along with the profile of species either present or absent in each supplement and used this information for dimensionality reduction and visualization with PCA (Figure 1 E). We observe no apparent clustering in this PCA plot based on marketed use for the probiotic. This result indicates that across all over-the-counter probiotic supplements, there is no specific combination of species that are consistently used to specifically support gut health, vaginal health, etc., in probiotic supplements. This result highlights the gap in understanding the functional capabilities of individual probiotic species. To elucidate the functional capabilities across probiotic species and how their functions compare to pathogenic and commensal species, we employed computational approach. Ultimately, this work will allow us to gain a deeper understanding of how probiotics can be used to positively impact human health, and is a step toward rational design of probiotic supplements and therapeutics.

### Metabolic network modeling of probiotic, commensal, and probiotic bacterial species

To assess functional capabilities of probiotic, commensal, and pathogenic bacterial species, we generated genome-scale metabolic network reconstructions (GENREs) representing 1,012 bacterial species using the automated GENRE construction tool, Reconstructor (Figure 2 A, Methods) (*33*). We created 779, 198, and 35 GENREs of commensal, pathogen, and probiotic metabolism respectively, which we call the CoPaPro collection. With these GENREs, we explored reaction content across probiotic, commensal, and pathogenic species.

**Figure 2.**
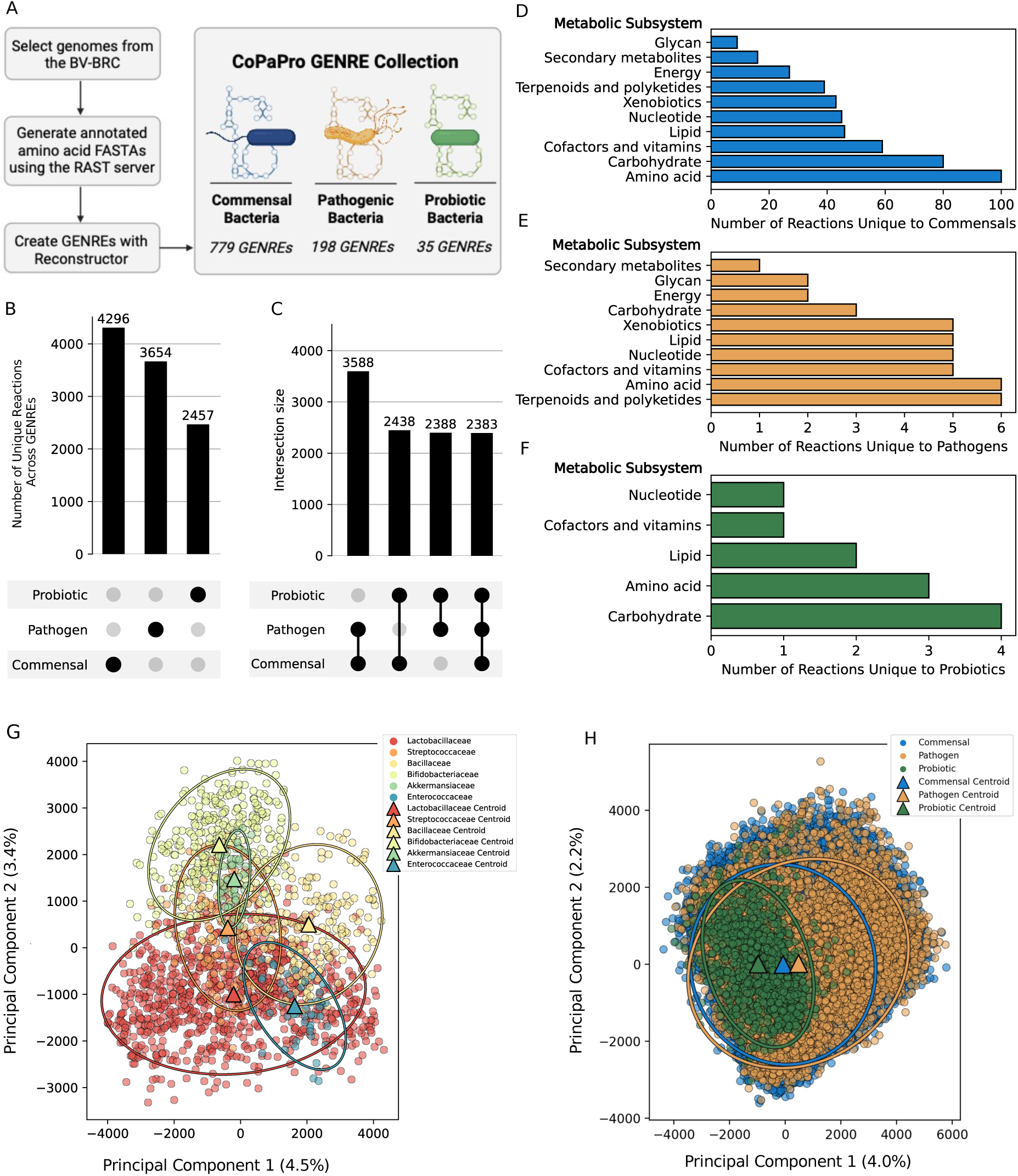
Description of genome-scale metabolic network models of probiotic, pathogen, and commensal metabolism. A) Workflow for generating the CoPaPro GENRE collection. B) Number of unique reactions across the GENREs of each category. C) Number of shared reactions between probiotic, pathogen, and commensal organisms. D-F) Metabolic subsystems of reactions that are unique to commensal (D), pathogen (E), and probiotic (F) GENREs. G) PCA of metabolic phenotypes of probiotic GENREs. Colors represent distinct taxonomic families. Triangles represent cluster centroid, ellipses plotted 2 standard deviations from the centroid. Kruskall-Wallis test for median distances between centroids, p-value = 1.16 e – 112. Cluster standard deviations are as follows: Lactobacillus: 995.19, Streptococcaceae: 533.53, Bacillaceae: 635.82, Bifidobacteriaceae: 609.72, Akkermansiaceae: 248.95, and Enterococcaceae: 488.13. H) PCA of metabolic phenotypes across probiotic, pathogen, and commensal GENREs. Triangles represent cluster centroid, ellipses plotted represent two standard deviations from the centroid. Kruskall-Wallis test for median distances between centroids, p-value = 0.

We observed that commensal species have the greatest number of unique reactions across GENREs (4,296 reactions) likely due to the size of the category and diversity of bacterial species it contains (Figure 2 B). On the other hand, probiotics have the fewest unique reactions (2,457 reactions), likely due to the small size of the category and the homogeneity of species it contains (*Lactobacillus, Bacillus, Bifidobacteria*). In the upset plot in Figure 2 C, we observe that there is significant overlap in reaction content between the three groups, with pathogens and commensals sharing the greatest number of reactions (3,588).

While we observed significant conservation in reaction content across the three groups, there were also reaction sets that were unique to each group. Across these unique reaction sets, there were differences in reaction metabolic subsystems. In commensal bacterial species, most of the unique commensal reactions are a part of the amino acid metabolism (Figure 2 D). Amino acid metabolism is an essential process for all organisms, but there have been studies that explicitly suggest that commensal bacteria are essential for the synthesis and absorption of certain amino acids like alanine, glutamate, and tryptophan (*34*, *35*). Alternatively, we observed that unique pathogenic reactions were mostly belonging to both the amino acid biosynthesis pathway, and the terpenoid and polyketide biosynthesis pathway (Figure 2 E). Many well-known antibiotics such as tetracycline are polyketides, most notably produced by *Streptomyces*, and some polyketides even act as virulence factors in pathogens (*36*, *37*). Finally, we observed that reactions unique to probiotic bacteria most often belonged to carbohydrate metabolism pathways (Figure 2 F). This result is consistent with previous studies that show carbohydrate metabolism is one key functions of probiotic bacteria, which aids digestion and promotes short chain fatty acid production, which are essential for fermenting dietary fiber and regulating inflammation (*38–41*). This presence of unique metabolic reactions and heterogeneity in the metabolic subsystems suggests that bacterial species from commensal, pathogen, and probiotic categories could occupy distinct metabolic niches (*42*).

### Understanding differences in metabolic niche through in silico simulation

We used the GENREs of commensal, pathogenic and probiotic metabolism from the CoPaPro collection to probe each species’ genotype-phenotype relationship using constraint-based reconstruction and analysis (COBRA). We generated metabolic phenotypes for each GENRE by generating flux distributions through flux balance analysis. Each generated flux distribution represents a distinct metabolic state of a specific organism. By sampling the flux solution space hundreds of times, we can gain an understanding of the range of all possible metabolic phenotypes for a given organism (*43*).

We first examined the range of metabolic phenotypes across all probiotic bacteria to gain a better understanding of functional similarities and differences within this category. To visualize flux distributions, we employed principal component analysis (Figure 2 G). We observed clustering of metabolic function in probiotics by family (p-value < 0.001). By examining similarities and differences in metabolic function in current probiotic species, we can select probiotic species that capture the widest range of metabolic functions to fill a wide range of metabolic niches. For example, combining species from the Lactobacillacea and Bifidobacteriacea families would cover a wide range of metabolic phenotypes. On the other hand, combining Akkermansiacea and Bifidobacteriacea species together may not have any added benefit than just administering Bifidobacteriacea strains alone.

Secondly, we were interested in gaining a deeper understanding of how commensal (blue), pathogenic (yellow), and probiotic (green) metabolic phenotypes are similar and different (Figure 2 H). Through this analysis, we observed many pathogenic and commensal metabolic phenotypes not captured by probiotic species. The gap in metabolic overlap we observe across current probiotics suggests that there is a large opportunity to expand species used in probiotic supplements. We will expand upon this analysis and identify ways to fill this metabolic gap in a specific use case; women’s health and the vaginal microbiome.

### Metabolic function in vaginal probiotic and commensal species

Through our survey of commercially available probiotics, we identified 23 supplements that claim vaginal health benefits containing 22 unique probiotic bacterial species with *Lactobacillus sp.* comprising 15 of the 22 unique species, likely because a *Lactobacillus* dominant vaginal microbiome is considered healthy (Figure 3 A) (*16*). However, certain populations do not exhibit *Lactobacillus* dominance nor symptoms consistent with vaginal dysbiosis (*20*) suggesting that non-Lactobacillus commensal members of the vaginal microbiome can be just as important for maintaining homeostasis and could be potential candidates for a novel probiotic consortium to promote vaginal health. To identify vaginal commensal species suitable for this role, we performed a combination of computational and *in vitro* analyses.

**Figure 3.**
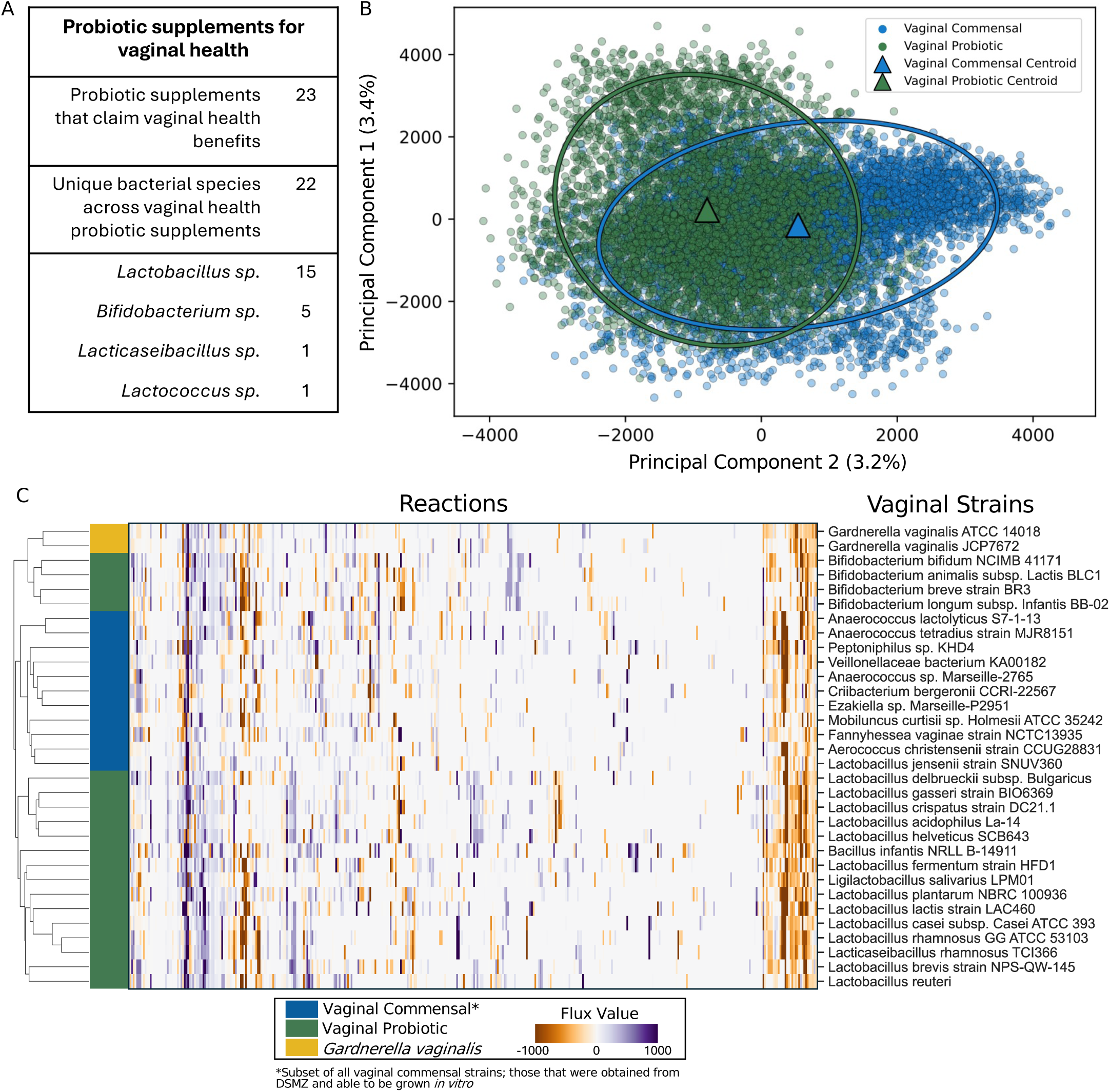
Functional analysis of vaginal commensal and probiotic species. A) Description of probiotic supplements that tout vaginal health benefits. B) PCA plot of metabolic phenotypes of vaginal commensal and probiotic species. Triangles represent cluster centroids, ellipses plotted 2 standard deviations from the centroid Kruskal-wallis test for median distances between centroids, p-value=2.2e-7. Vaginal commensal standard deviation: 897.7. Vaginal probiotic standard deviation: 863.35. C) Hierarchical clustering of metabolic phenotypes across vaginal probiotic species, two *G. vaginalis* strains, and culturable vaginal commensal species. Metabolic phenotype vectors are calculated as the per-reaction median value across 500 flux samples.

We began by identifying vaginal commensal strains and vaginal probiotic species from the larger CoPaPro collection and simulated metabolic phenotypes using flux balance analysis. Through dimensionality reduction and visualization of the metabolic phenotypes, we observe that there are significant differences in metabolism between species used as vaginal probiotics in commercially available supplements, and the native flora of the vaginal microbiome, with the commensal species covering a wider range of metabolic functions (vaginal commensal standard deviation: 897.7; vaginal probiotic standard deviation: 863.35; Kruskal-Wallis test for median distances between centroids: p=2.2e-7) (Figure 3 B). Consistent with what we observed in Figure 2 H, there are metabolic functions of vaginal commensals that are not captured with the current suite of vaginal probiotic species. This result presents an opportunity to leverage native commensal members of the vaginal microbiome as possible novel vaginal probiotic strains.

Prevention of *G. vaginalis* growth, frequently associated with BV, is one specific target we could consider when designing next-generation probiotics for the vaginal microbiome (*44*) through identifying commensal vaginal species with a metabolic niche most like *G. vaginalis*, thereby preventing proliferation through resource competition. To identify these species, we began by examining the metabolic profiles of two *G. vaginalis* strains, commensal vaginal species that can be grown *in vitro* (necessary for downstream *in vitro* assays), and vaginal probiotic species, to identify the median metabolic signature of each strain. Then, using hierarchical clustering, we can examine the similarities and differences in metabolic signatures across all strains (Fig 3 C).

Ultimately, we observe that *G. vaginalis* strains have very similar functional signatures to *Bifidobacterium* species that are currently used in vaginal probiotics. Interestingly, the *Lactobacillus* and *Bacillus* species used in vaginal probiotics are more functionally like commensal vaginal species than *G. vaginalis.* This result implies that overlap in metabolic niches may not be the only reason for an effective probiotic. According to these *in silico* results, *Lactobacilli* do not have similar metabolic profile to *G.vaginalis*, but we know that *Lactobacilli* are essential for maintaining vaginal health through secretion of lactic acid to maintain a low pH environment. This result could suggest that when considering the rational design of probiotics, preventing *G. vaginalis* colonization may not be a function of only overlapping niche or potential for competition, but may be a more nuanced problem which also considers additional environmental factors. To gain more insight into the balance between resource competition and environment factors driving the prevention of *G. vaginalis* colonization, we performed an *in vitro* spent media assay to model competition between *G. vaginalis* and vaginal commensal species.

### Spent media assay reveals commensal vaginal species whose spent media inhibits G. vaginalis growth

Our computational metabolic phenotype analysis revealed that there are distinct functional groups of vaginal commensal species. We observed that some vaginal commensals have similar metabolic phenotypes to *G. vaginalis* strains, implying these species could exhibit competitive interactions. Vaginal commensal species that compete with *G. vaginalis* could be key candidates for a novel probiotic consortium preventing *G. vaginalis* colonization. To further understand differences in metabolic functionality across vaginal commensals and identify which commensals effectively inhibit *G. vaginalis* growth, we designed an *in vitro* spent media assay. This assay is a proxy for assessing competition between vaginal commensals and *G. vaginalis*; allowing us to determine which vaginal commensal bacteria produce metabolic byproducts that would inhibit the growth of *G. vaginalis.* Through this experiment, we can better understand if a stable population of the vaginal commensal could prevent the proliferation of a *G. vaginalis* population. We began by generating 11 spent media conditions, one for each of the culturable commensal vaginal species shown in Figure 3 C, that were able to be cultured *in vitro*. Then, an overnight culture of *G. vaginalis* was inoculated into each species’ spent media and growth was monitored through stationary phase (Methods).

We observed some variation in the *G. vaginalis* growth profiles on the 12 media conditions (PGY (Modified) control, and 11 spent media conditions) (Figure 4 A). More specifically, we see three distinct response groups; Non-inhibitory (N), Moderate (M), and Inhibitory (I) (pairwise t-tests of area under the mean growth curve: N vs M: p-value < 0.01, N vs I: p-value < 0.001, and I vs M: p-value < 0.001) (Figure 4 B). *G. vaginalis* grown on non-inhibitory media (*Mobiliuncus curtisii, Veillonellaceae bacterium, and Ezakiella massiliensis* spent media) had the same growth dynamics as *G. vaginalis* grown on the PGY (Modified) media control. In the moderate group, we see a significant change in the area under the growth curve of *G. vaginalis* grown in *Peptoniphilus vaginalis, Anaerococcus marseille, Criibacterium bergeronii,* and *Aerococcus christensenii* spent media compared to the uninhibited group (p-value < 0.01). In the inhibited group, we see a statistically significant difference in the area under the growth curve of *Gardnerella vaginalis* grown in *Fannyhessea vaginae* spent media*, Lactobacillus jensenii, Anaerococcus lactolyticus,* and *Anaerococcus tetradius* when compared to the uninhibited (p-value < 0.001) and moderate (p-value <0.001) conditions.

**Figure 4.**
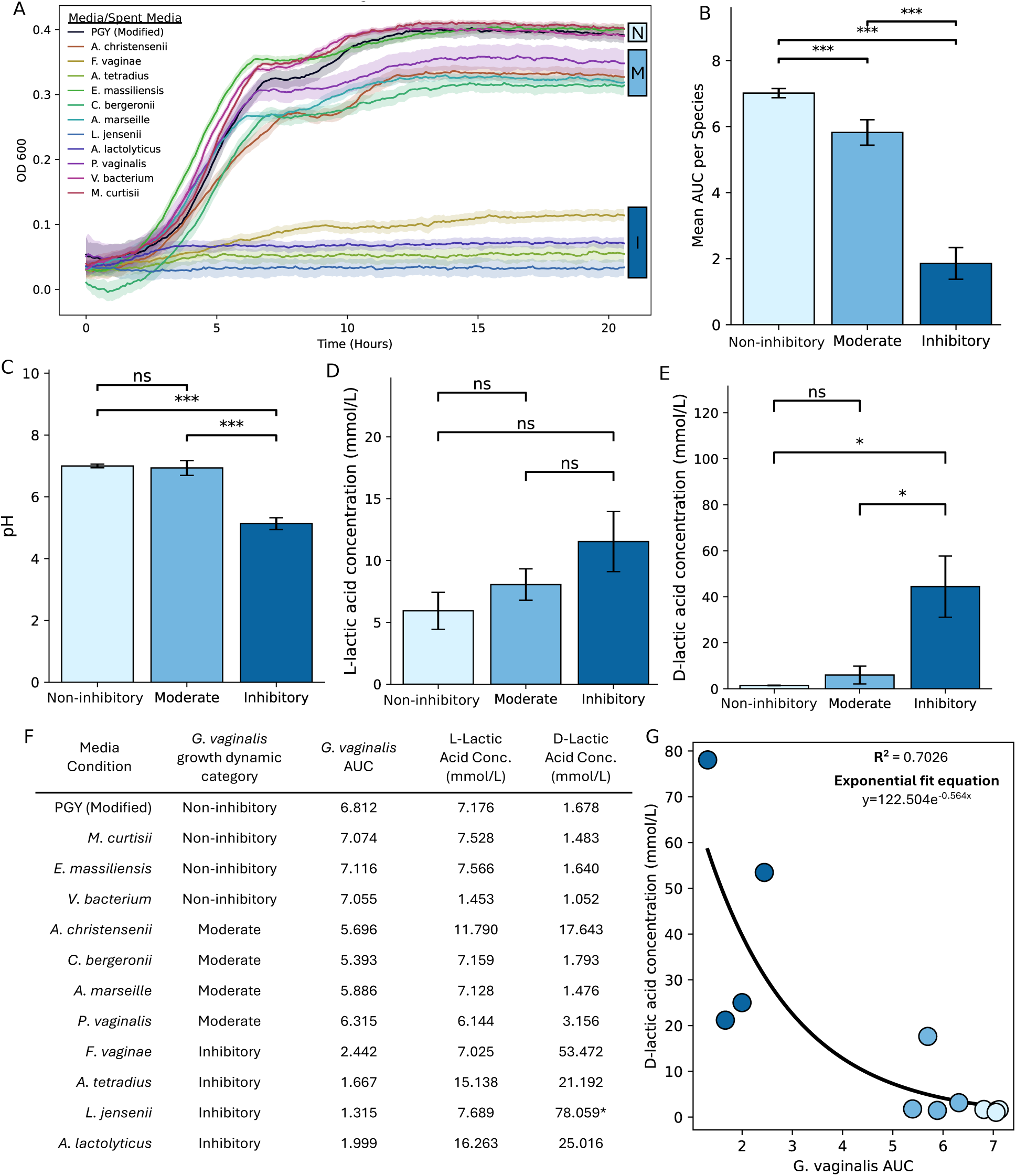
*In vitro* spent media assay and lactic acid quantification. A) Growth curves of *G. vaginalis* 14018 in 12 media conditions (11 spent media and PGY (Modified) control). Three growth dynamics are specified to the right of the growth curves. N: non-inhibitory, M: moderate, I: inhibitory. B) Quantification of area under the growth curve for each of the three growth dynamic categories. Pairwise t-tests, one tailed: N vs M: p-value = 5.68e-4, n vs I: p-value = 4.19e-7, M vs I: p-value = 6.76e-6. C) pH of the spent media in each growth dynamic category. Pairwise t-tests, one tailed: I vs M: p-value = 6.46e-4, I vs N: p-value = 6.32e-4, M vs N: p-value = 0.4039. D) L-lactic acid concentration in the spent media of each growth dynamic category. Pairwise t-tests, one tailed: I vs M: p-value = 0.133, I vs N: p-value = 0.0532, M vs N: p-value = 0.1605. E) L-lactic acid concentration in the spent media of each growth dynamic category. Pairwise t-tests, one tailed: I vs M: p-value = 0.0292, I vs N: p-value = 0.02418, M vs N: p-value = 0.1632. F) Concentration of L-lactic acid, D-lactic acid, area under the *G. vaginalis* growth curve, and growth dynamic category of each spent media condition. Asterix (*) indicates that this concentration was beyond the standard curve and was extrapolated. G) Relationship between D-lactic acid and *G. vaginalis* AUC. R^2^=0.7026, exponential fit equation = y=122.504e^-.564x^. Bar chart significance key: *** = p < 0.001, ** = p < 0.1, * = p < .1, ns = not significant.

The three distinct patterns of *G. vaginalis* growth dynamics in the 11 spent media conditions suggests that there could be multiple mechanisms of *G. vaginalis* growth inhibition at play. We next wanted to determine if resource scarcity and competition or environmental factors were driving *G. vaginalis* growth inhibition in the inhibitory spent media.

### In vitro assays uncover D-lactic acid as major driver of G. vaginalis growth inhibition

As mentioned previously, it is well-known that the *Lactobacillus* dominance in the vaginal microbiome is responsible for protecting against pathogen invasion by maintaining an acidic environment through lactic acid production (*16*). Because of this, we first determined if the differences in growth dynamics of *G. vaginalis* observed in the different spent media conditions were influenced by pH of the spent media.

We observed significant differences in pH across the spent media of the three growth dynamics groups (N, M, and I) (Figure 4 C). The spent media that inhibited *G. vaginalis* growth had a significantly lower pH than the spent media of the moderate and uninhibited groups (I vs N: p-value<0.01, I vs M: p-value<0.01). However, there was no significant difference between the pH in the uninhibited and moderate groups (N vs M: p-value > 0.8). This result suggests that pH of the spent media plays a major role in inhibiting the growth of *G. vaginalis*. To determine if the low pH of the inhibitory spent media is due to lactic acid production, we performed both D-lactic acid and L-lactic acid assays (Methods).

The mmol/L concentrations of L-lactic acid and D-lactic acid determined by the assays are reported in Fig 4 F. Ultimately, we observed no significant difference in the concentration of L-lactic acid between the inhibitory, non-inhibitory, and moderate spent media (Figure 4 D). However, the inhibitory spent media contained more L-lactic acid on average than the moderate and non-inhibitory spent media, and the moderate spent media contained more L-lactic acid on average than the non-inhibitory spent media (Figure 4 D). We did see significant differences in D-lactic acid concentration in the inhibitory spent media condition compared to the moderate and non-inhibitory spent media conditions (Figure 4 E). Additionally, we observed that there is a relationship between the *G. vaginalis* area under the curve (AUC) and D-lactic acid concentration of the spent media (Figure 4 G). Taken together, these results suggest that D-lactic acid production is a major driver for *G. vaginalis* growth inhibition in the inhibitory spent media condition.

These results are consistent with the general knowledge that *Lactobacillus* species are typically present in a healthy vagina due to their ability to produce lactic acid and lower vaginal pH. However, we observed significantly high levels of D-lactic acid produced by *F. vaginae*, *A. tetradius*, and *A. lactolyticus*, despite not being *Lactobacillus* species. These results suggest that any bacteria that produces high levels of D-lactic acid could be just as effective as *Lactobacillus* at preventing *G. vaginalis* colonization.

While these results help to elucidate that D-lactic acid is important for *G. vaginalis* growth inhibition, further studies would need to be performed to elucidate the driver of the differences in *G. vaginalis* growth dynamics observed between the non-inhibitory and moderate spent media groups. Based on preliminary computational analyses presented in S4, there is evidence that metabolic resource competition between commensals and *G. vaginalis* (resource competition) may be a driver of these observed differences in growth dynamics. In this analysis we observed some clustering of metabolic phenotypes of moderately inhibitory and inhibitory vaginal commensals with *G. vaginalis* strains suggesting metabolic niche overlap, however this analysis would need to be expanded and refined to better define the potential competitive mechanisms of inhibition.

## Discussion

There has been extensive research on the positive health outcomes linked to probiotic intake, resulting in hundreds of probiotic supplement products being available to consumers over-the-counter (*3–6*). However, we lack a mechanistic understanding of the bacterial species contained within these supplements and how they can support health of the system of interest. Consequently, we explored the range of metabolic functions across species in over-the-counter probiotic supplements using a metabolic network modeling approach to identify potential opportunities to expand species used in current supplements to capture a broader range of metabolic functions.

We observed that the metabolic functions captured across probiotic species used in supplements was only a small subset of the functions captured across all pathogenic and commensal species. Furthermore, we suggest that metabolic network modeling could play a part in rationally designing combinations of probiotic bacteria to add to a supplement to maximize the range of metabolic functions captured. In the future, if sequences of the individual proprietary strains used in probiotic supplements were made available, we could expand this analysis to be strain-specific. Additionally, it would be valuable to understand how species in each supplement function synergistically to determine if combining species could expand metabolic functionality. Furthermore, colonization rates of probiotic species are generally low and quite variable (*45*). Consequently, considering competitive and mutualistic interactions between probiotic species and the native microbes should be explored in the future. Using a community modeling approach could help us better identify gaps in probiotic function and how these gaps could be filled with novel probiotic communities rather than individual strains.

Additionally, we focused on the vaginal microbiome to identify gaps in probiotics used to support vaginal health. We observed that most vaginal microbiome-related probiotics consist of lactic acid bacteria like *Lactobacilli* which are known to support vaginal health by maintaining an acidic vaginal pH. In this work we wanted to identify commensal microbes in the native vaginal flora that prevent *G. vaginalis* growth, which could be candidates for novel vaginal probiotic species. Through spent media experiments to simulate resource competition between *G. vaginalis* and a commensal vaginal species, we identified four commensal vaginal species (*A. lactolyticus, A. tetradius, L. jensenii,* and *F. vaginae*) that inhibit *G. vaginalis* growth through D-lactic acid production. However, D-lactic acid production could depend on the media context; in this study we only used PGY-mod (Methods) to create our spent media conditions. This study could be repeated using spent media generated on media that better mimics the vaginal environment like NYCIII or synthetic vaginal media (SVM). Additionally, we could utilize a co-culture assay to examine competition and growth inhibition in real time, to see how the dynamics of *G. vaginalis* are altered in the presence of the vaginal commensal. Additionally, there could be an opportunity to explore how *G. vaginalis* would grow on the spent media of a vaginal commensal community. This would provide an opportunity to see if combinations of the commensal vaginal microbes may be able to more strongly inhibit *G. vaginalis* growth than individuals. Furthermore, this analysis could be expanded to other microbiomes to identify commensals that serve as competitors for certain infectious pathogens. Overall, we believe this metabolic network modeling approach combined with *in vitro* experimentation is a promising step toward rationally designed probiotics for targeted uses, specifically for supporting vaginal microbiome health through *G. vaginalis* growth inhibition.

There should not be a one-size fits-all approach to treating bacterial vaginosis; there is a need for patient-specific therapies that can consider variations in vaginal microbiome composition. Specifically, post-menopausal women have been shown to have significantly different vaginal microbiome compositions in both healthy and dysbiotic states due to differences likely due to differences in systemic estrogen (*46*). Most BV diagnostics and therapeutics have been designed with only reproductive-age women in mind, which could render these therapeutics less effective for post-menopausal women with BV. The number of post-menopausal women (55+) presenting with vaginitis-associated symptoms in the clinic is not a small portion. According to TriNetX, Of 35,040 reported vaginitis cases in the UVA health system, 7,580 (22%) of these cases were in women > 55 years old (S5, Methods). This emphasizes the need for vaginitis/BV treatment options specifically for post-menopausal women. The systems biology approach presented here could extended to identify possible probiotic candidates to protect against non-*Gardnerella* dominant BV, which would lend a more patient-specific therapeutic approach.

## Author Contributions Statement

E.G. wrote initial manuscript draft

E.G. performed computational studies

E.G. and G.K. performed experimental analyses

J.P. and G.K. supervised the work

E.G., G.K., and J.P., edited and approved of the final manuscript

## Data and Materials Availability

All raw data, scripts, and metabolic network models used in this study are available online: https://github.com/emmamglass/ProbioticsAndWomensHealth

## List of Supplementary Materials

Methods

S1-S8

## Methods

### Survey of Commercial Probiotics

We performed a survey of probiotics available for purchase on the CVS, Walgreens, and Walmart websites, the top three pharmacies in the USA by number of locations (9,554, 9,398, and 6,860 stores respectively) (*29*). In this survey, we collected the name of each probiotic supplement product, specified uses (e.g., gut, vaginal, urinary), and all probiotic strains included in each supplement. Additionally, we determined the total number of strains and the total number of species present in each probiotic supplement. We then created a binary species presence dataframe (row: probiotic supplement product, column: probiotic species, 1: species present, 0: species absent). We then used this dataframe for dimensionality reduction and visualization with PCA (sklearn in Python) to observe clustering patterns (*47*).

### Metabolic Network Reconstruction of Commensal, Pathogenic, and Probiotic Species

We generated 1,007 genome scale metabolic network reconstructions (GENREs) of commensal, pathogenic, and probiotic bacterial species. To do this, we began by selecting genome sequences from the BV-BRC (*48*). We selected one genome sequence for each of the 35 bacterial probiotic species identified in our survey of commercially available probiotics. The criteria for selecting probiotic genome sequences were as follows: 1) Select the representative or reference sequence of a given species if it exits, 2) If there is no representative or reference sequence, randomly select a sequence that is considered good quality and complete. Of the 35 probiotic sequences selected, 26 were considered reference or representative sequences, and 9 sequences were considered good and complete without the reference or representative designation. We selected pathogen sequences in a similar manner. Using the previously published database of metabolic network models as a guide, we selected all reference and representative sequences of the species included in the PATHGENN database (*42*), resulting in 197 pathogenic sequences selected. Finally, we considered commensal bacteria to be any species in the BV-BRC database that was not considered pathogenic or probiotic. We selected all reference and representative commensal sequences that met these criteria, resulting in 775 commensal sequences.

After selecting the sequences, we generated an annotated protein sequence through an automated pipeline. We used this annotated protein sequence as an input to Reconstructor, a tool used for automated GENRE creation (*33*). We used Reconstructor to generate genome scale metabolic network reconstructions of all 1,007 sequences. All GENREs were created in the context of the same rich media.

### Reaction and subsystem analysis

We identified reactions that were unique to commensal, pathogen, and probiotic bacterial species through model analysis and visualized this data using an upset plot (Figure 2 B, 2 C). We then took the list of unique metabolic reactions from each group (commensal, pathogenic, probiotic) and identified the metabolic subsystem to which they belong. We did this by querying the KEGG API to identify the general subsystem that each of these unique reactions belongs to (*49*). We then displayed the number of unique reactions in each group that were a part of each metabolic subsystem.

### Flux sampling and metabolic phenotype analysis

We used the Gapsplit algorithm for sampling flux distributions (*50*). We generated 500 flux distributions for each GENRE to most accurately capture the range of all metabolic functions (metabolic phenotypes). To visualize similarities and differences in flux distributions across commensal, pathogen, and probiotic species, we used principal component analysis (PCA) through the sklearn python package (*47*). We reduced each flux distribution to a two-dimensional space (two principal components) to plot. In Figure 2 H, we plotted commensal bacterial species (bottom), then pathogenic (middle), then probiotics (top) to best visualize the metabolic phenotype coverage of group. We used the same method of flux sampling and dimensionality reduction in Figures 2 G and 3 B.

### Heatmap and hierarchical clustering of metabolic flux through G. vaginalis, vaginal commensal, and vaginal probiotic bacterial species

We generated 500 flux distributions per model of interest using Gapsplit (*50*). Then, we generated a median flux vector across all 500 samples per each model. We then combined all median flux vectors across models of interest into one data frame, removed low variance reactions (variance threshold of 0.1), and dropped highly correlated reactions (correlation threshold of 0.9). After these pre-processing steps, we generated a cluster map using seaborn (*51*), clustering on both rows (species-specific models) and columns (reactions) using the Canberra distance. This method was applied to the heatmaps in Figures 3 C and S4.

### In vitro spent media assays

We obtained 16 vaginal commensal isolates from the DSMZ (strain names in S6). We selected these isolates from the larger collection of identified vaginal commensal organisms for several reasons: 1) the isolates were easily obtainable from one reputable source, 2) we selected strains that were specifically isolated from the human vagina. There were strains of some isolates that were available but were not isolated from the vagina; we did not select these.

Ultimately, we were able to successfully grow 11 of these species robustly in anaerobic conditions. After successfully growing these bacterial species in their specified media, we determined a media condition that would successfully grow these 11 species as well as *Gardnerella vaginalis.* We determined the media that was most successful at robust growth was PGY media (DSMZ) modified with HEPES (6.3 mL) and 10% FBS. We also tried unmodified PGY, NYCIII, and NYCIII media modified with Vitamin K (100uL) and Hemin (5mL) (S7).

After determining a universal media condition (PGY modified; PGY-mod that would lead to maximal growth across isolates, we next needed to collect spent media. We inoculated each isolate into 10 mL of PGY-mod. For each isolate, we repeated this 5 times. Each 10mL aliquot of spent media was pooled to generate 50 mL of spent media per isolate.

Then, to determine if *G. vaginalis* would grow on the spent media of each isolate, we performed a spent media assay. This assay involved inoculating *G. vaginalis* into 5mL of PGY-mod and was allowed to grow for 24 hours. After 24 hours, the culture was spun down at 6500 rpm for 5 minutes, media was aspirated, and culture was resuspended in fresh PGY-mod media. Then, *G. vaginalis* was inoculated into the spent media in a 12 well plate at an OD of 0.1 and allowed to grow until stationary phase with continuous growth monitoring using a Cerrillo stratus plate reader. We repeated this assay with two different strains of *G. vaginalis*; GV14018 (results show in Figure 4 A) and JCP7672 (S6) to ensure the trends we observed were consistent across strains. We observed inhibition of *G. vaginalis* JCP767 in *A. tetradius, A. lactolyticus, L. jensenii,* and *F. vaginae* spent media as well. We observed growth of JCP7672 in the moderate and Non-inhibitory spent media that were mostly consistent with GV14018, but there was some variation (S8).

### Area under the growth curve calculations and identifying differences in growth dynamics

We calculated area under each growth curve in Figure 4 B using the simps integration method from the scipy package (*52*). We used the mean value across replicates for this growth curve calculation. To determine statistical difference between the three observed groups of growth curves, we used a one-way ANOVA test, with subsequent pair-wise t-tests between groups, with a significance value of 0.05.

### Spent media pH measurements and differences between groups

We used a pH meter (Accumet basic AB15 pH meter) to determine the pH of each media condition used in the spent media assay with accuracy to 0.01. Then, to determine statistically significant differences in pH between the three growth dynamics groups we used pairwise Welch’s t-tests, with a significance value of 0.05.

### L-lactic acid and D-lactic acid assays

We quantified concentrations of D-lactic acid and L-lactic acid using the Novus Biologicals D-Lactic Acid/Lactate Assay Kit (Colorimetric) NBP3-25788 and L-lactic Acid Assay Kit (Colorimetric) NBP3-25875 respectively. After completing the specified kit protocol, OD was read at 530nm using a TECAN plate reader. The assay was re-run (standards and samples) at dilution for samples that read above the standard curve range. D-lactic acid and L-lactic acid concentrations were computed according to the standard curve.

### TriNetX cohot exploration

To explore the number of vaginitis cases presented in the UVA health system, we used TriNetX. The data used in this study was collected on June 5, 2025 from the TriNetX University of Virginia Network, which provided access to electronic medical records (diagnoses, procedures, medications, laboratory values, genomic information) from 730,801 patients from health care organization. This retrospective study is exempt from informed consent. The data reviewed is a secondary analysis of existing data, does not involve intervention or interaction with human subjects, and is de-identified per the de-identification standard defined in Section §164.514(a) of the HIPAA Privacy rule. The process by which the data is de-identified is attested to through a formal determination by a qualified expert as defined in section §164.514(b)(1) of the HIPAA Privacy Rule. This formal determination by a qualified expert refreshed on December 2020.

---

**S1.**
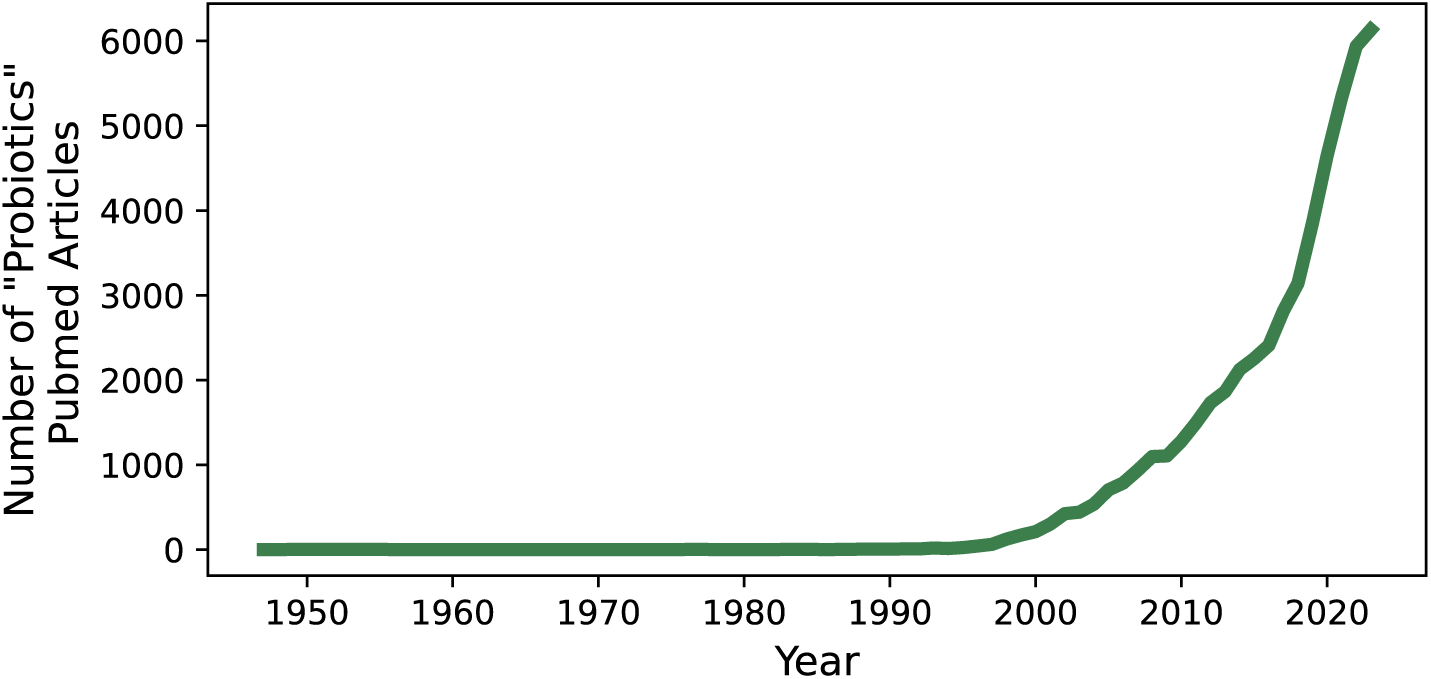
Number of “Probiotics” PubMed articles published each year since before 1950.

**S2.**
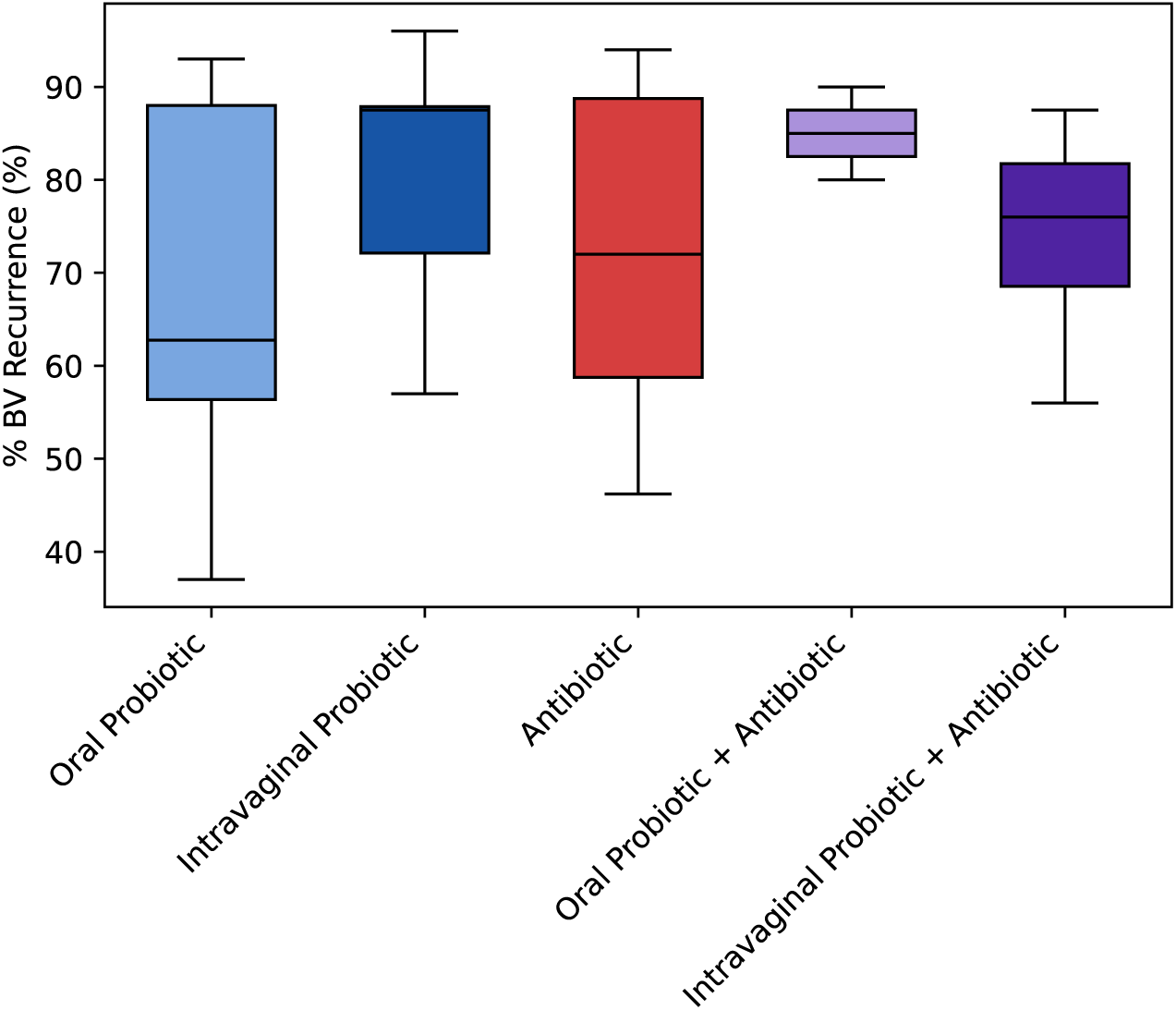
Reported BV recurrence rates after probiotic and antibiotic treatments. This data was compiled from 32 independent clinical studies. Oral probiotic: n = 8, Intravaginal probiotic: n = 6, Antibiotic: n = 25, Oral Probiotic + Antibiotic: n = 2, Intravaginal Probiotic + Antibiotic: n = 7.

## S3 - Bibliography of Studies Used in Meta-analysis

**S4.**
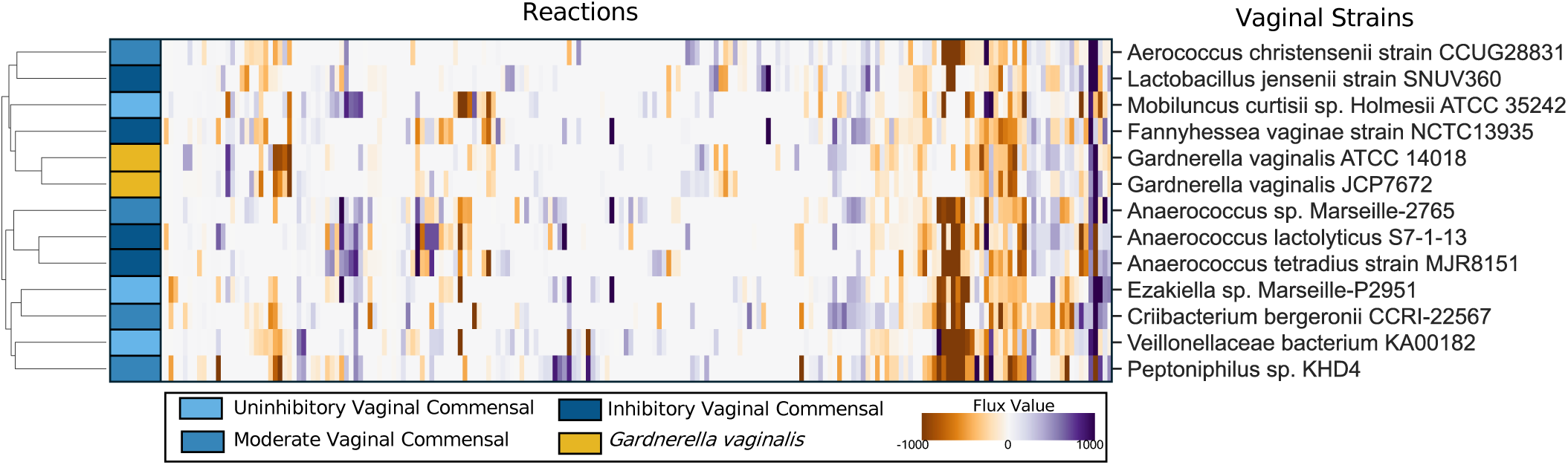
*In silico* metabolic phenotype comparison between vaginal commensals and *G. vaginalis* strains. Hierarchical clustering of metabolic phenotypes across two *G. vaginalis* strains, and culturable vaginal commensal species used in the in vitro spent media experiment, rows are colored by *G. vaginalis* growth dynamic category. Metabolic phenotype vectors are calculated as the per-reaction median value across 500 flux samples.

**S5.**
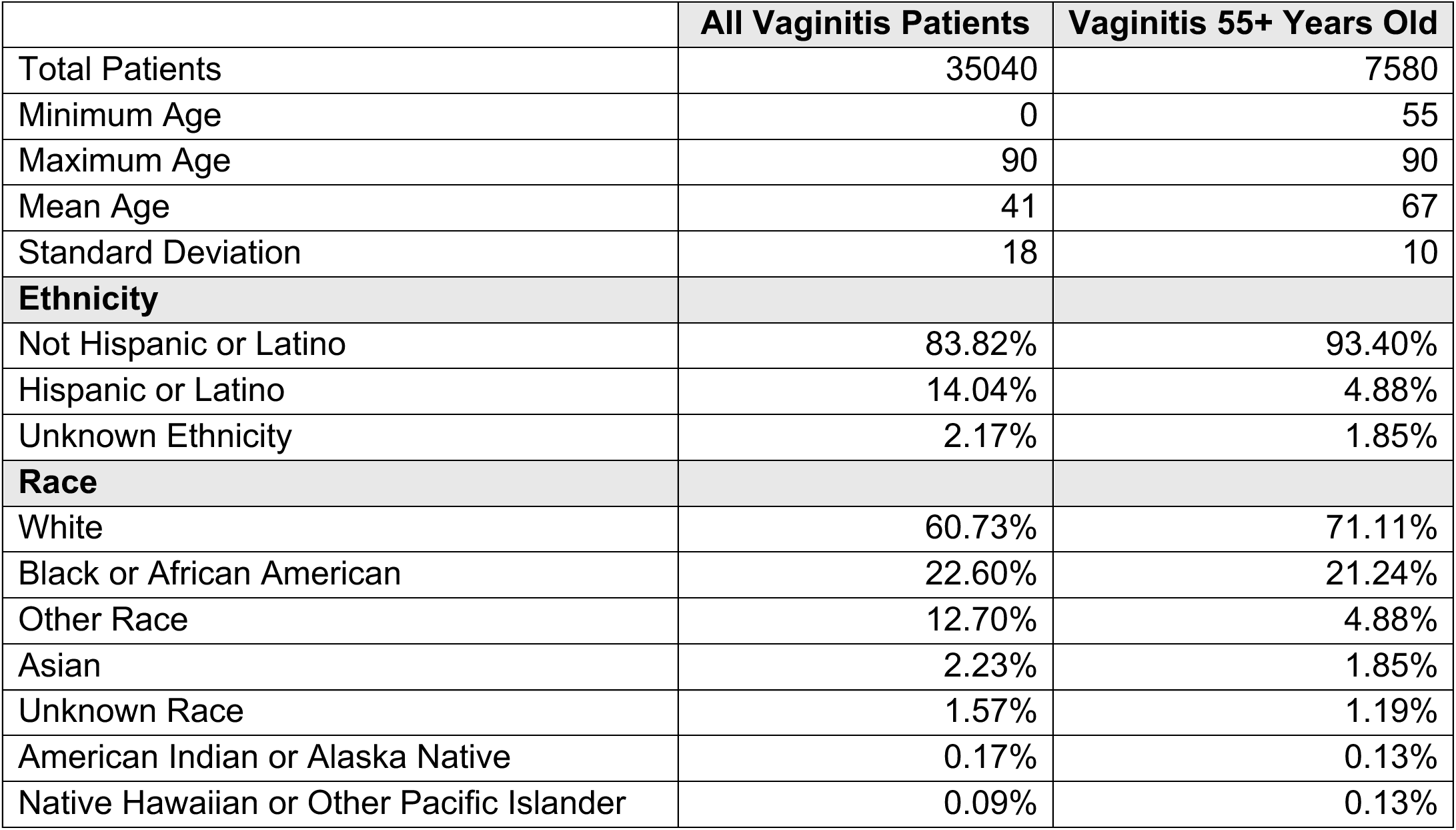
Patients presenting with vaginitis associated symptoms in the in the UVA health system.

**S6.**
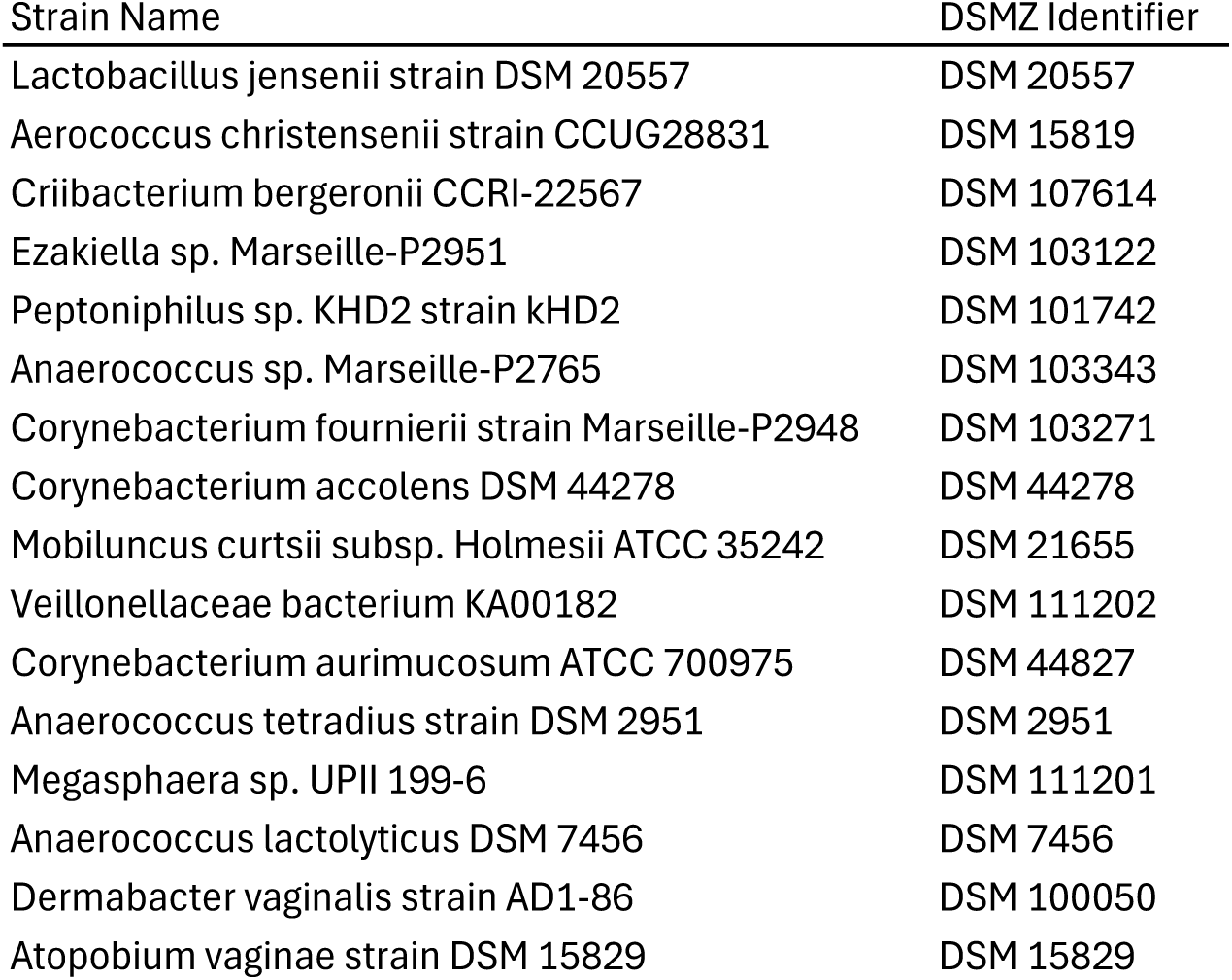
List of vaginal commensal isolates obtained from the DSMZ.

**S7.**
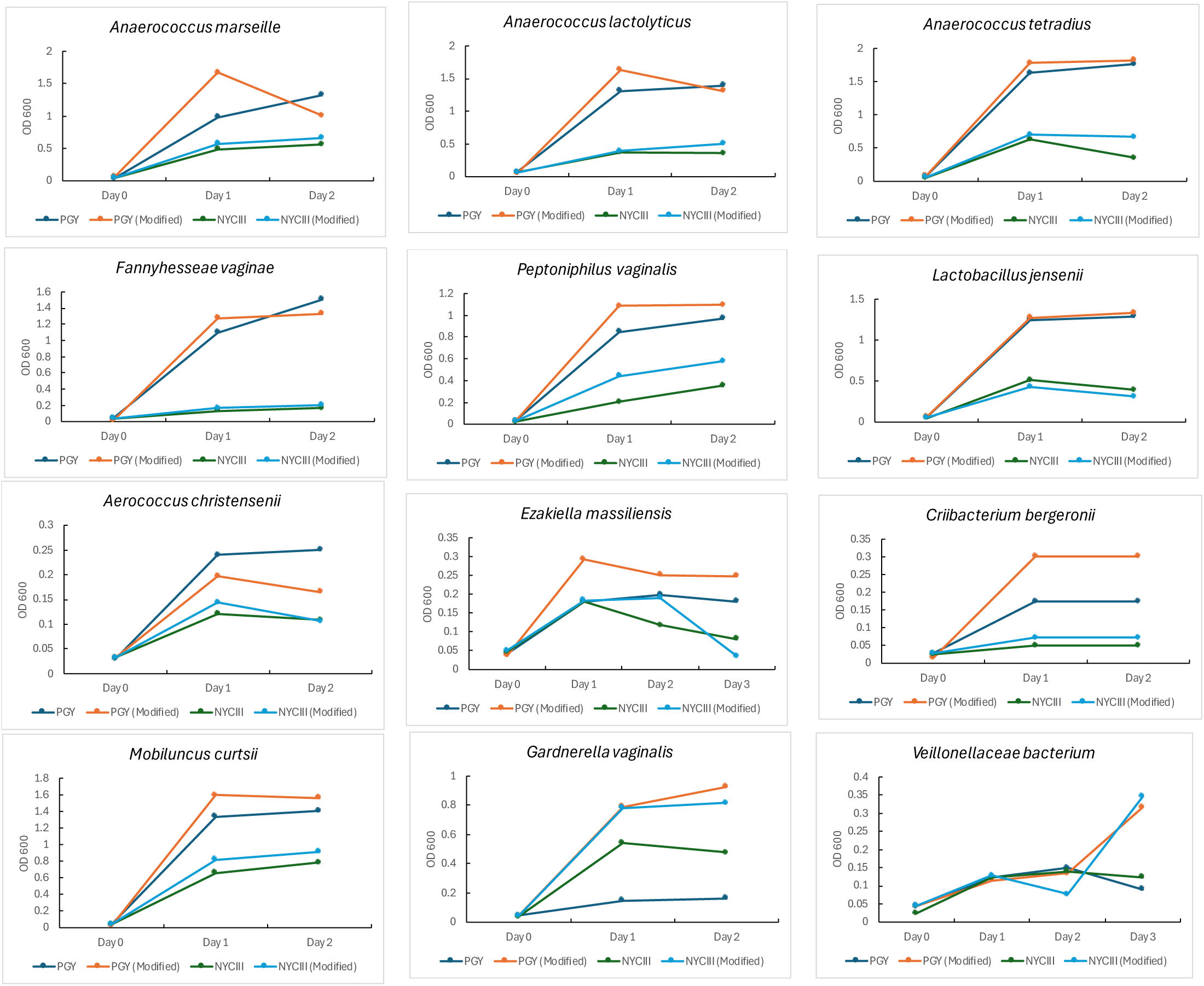
Comparison of culturable vaginal commensal species growth in four media conditions. Growth over 48 and 72 hour periods in PGY, PGY (Modified), NYCIII, and NYCIII (Modified).

**S8.**
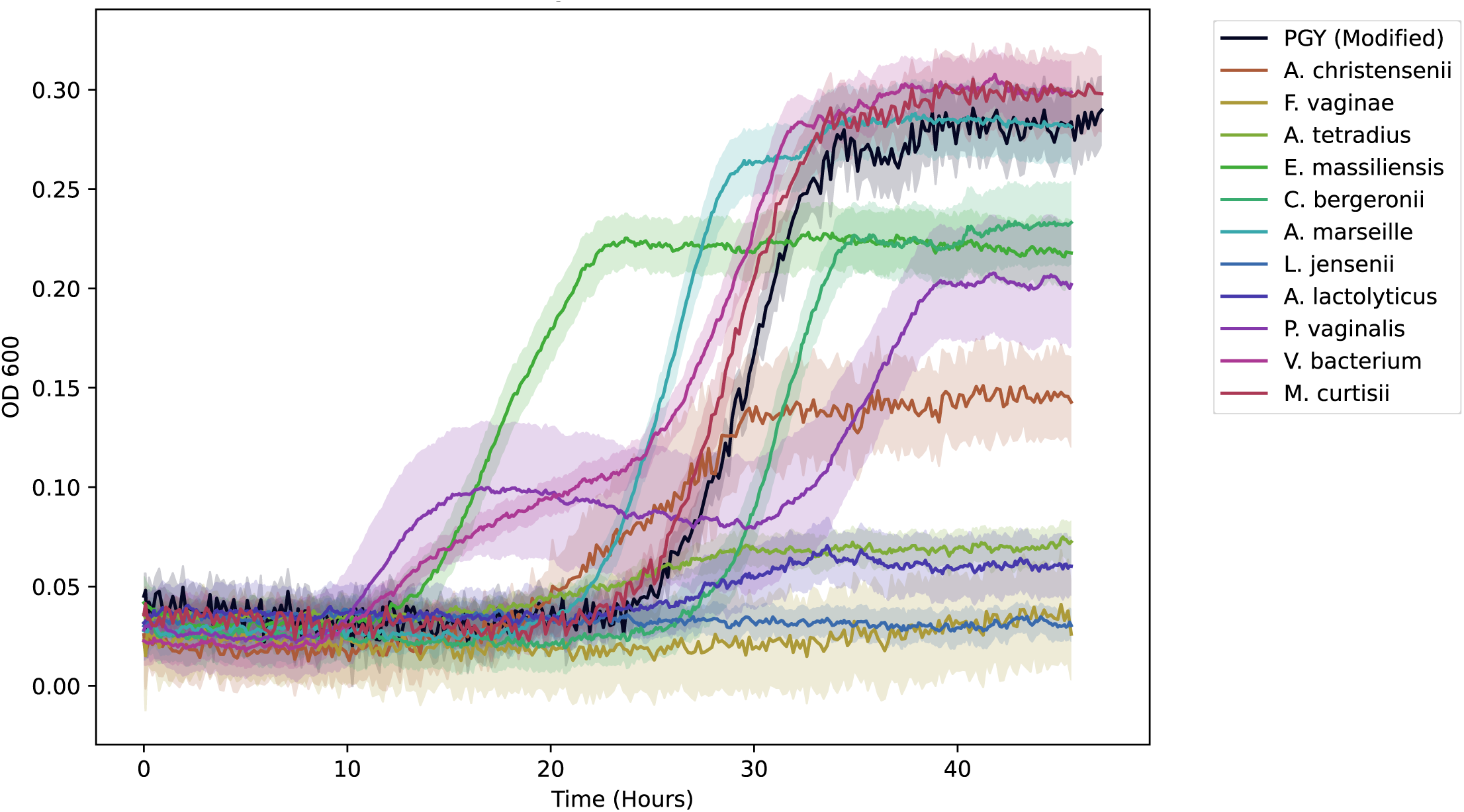
Growth of *G. vaginalis* JCP7672. This alternative *G. vaginalis* strain was grown on 11 spent media conditions and PGY (modified) control.

